# Pressure-induced membrane tension mechanically opens the germinant receptor GerA ion channel to trigger bacterial spore germination

**DOI:** 10.64898/2026.08.18.745453

**Authors:** Lei Rao, Tianyu Zhang, Ziqi Gong, Ke Liu, Yonggang Wang, Bing Zhou, Yongqiang Gao, Peter Setlow, Xiaojun Liao

## Abstract

High pressure (HP) can trigger bacterial spore germination, acting either through germinant receptors (GRs) or the SpoVA channel. However, the mechanism by which HP activates these membrane-embedded proteins remains elusive. Here, using *Bacillus subtilis*, we demonstrate that the GerA germinant receptor (GR) is the primary target of moderate HP (50-300 MPa). Mutagenesis reveals that pore-lining residues within the GerA ion channel are essential for the pressure response, whereas canonical ligand-binding and intramembrane signaling residues are dispensable. We then propose a <u>s</u>tretch-to-<u>o</u>pen (STO) model, in which HP differentially compresses the more compliant inner membrane (IM) relative to the rigid spore core, generating lateral membrane tension that promotes opening of the GerA channel. *In situ* membrane tension measurements indicate HP-induced compression of IM phospholipids and elevated membrane tension. This tension-dependent gating is further supported by the pressure-dependent phenotypic rescue of GerA channel mutants. Consistently, HP increases IM permeability to water-soluble and membrane-impermeable agents (propidium iodide and formaldehyde), an effect potentiated by GerA, indicating concomitant opening of GerA by HP. Furthermore, modulating IM fluidity via heat activation or decoating altered membrane physical properties and delayed HP-induced germination, establishing the IM as the critical mechanical transducer. Additionally, computational modeling and calculations support faster compression of the IM than of the core under HP, rationalizing the source of tensile stress. Together, our findings establish a novel mechanism of HP-induced GerA activation via the STO model: HP compresses the IM, generates lateral tension, and promotes opening of the GerA ion channel to trigger bacterial spore germination.

**Importance:** Bacterial spores are notoriously resilient, posing significant challenges to health and industry. While high pressure is known to trigger spore germination, the biophysical mechanism has remained elusive. Here, we provide evidence that high pressure mechanically activates the germinant receptor GerA via a “stretch-to-open” mechanism. Because the inner membrane is highly compressible compared to the rigid spore core, pressure induces membrane compression that generates lateral tension. This tension promotes opening of the GerA ion channel. Our study identifies the inner membrane as a critical mechanical transducer, addressing a long-standing question in spore biology and suggesting a distinct mechanosensory adaptation involved in spore revival.

## Introduction

Bacteria of the orders *Bacillales* and *Clostridiales* can differentiate into dormant, resistant spores under nutrient limitation, a state that confers remarkable resilience against heat, chemicals, radiation, and desiccation (1–4). Due to their resilience, spores persist widely in natural and man-made environments, with significant implications for health and industry (1, 5). Pathogenic species such as *Bacillus anthracis*, *B. cereus*, and *Clostridium botulinum* pose direct threats through toxin production (6, 7), while spoilage organisms such as *Geobacillus stearothermophilus* and *Alicyclobacillus acidoterrestris* withstand routine food-processing treatments, leading to economic losses (4, 8, 9). In their dormant form, spores are metabolically inactive and relatively benign; it is only upon germination and return to vegetative growth that they regain metabolic activity and become hazardous. Consequently, germination, which is the critical first step in spore revival, represents a key control point for both spore biology and applied microbiology (10–14).

In *Bacillus* species, germination is typically initiated when small-molecule germinants, such as amino acids or sugars, bind to specific germinant receptors (GRs) located in the spore’s inner membrane (IM) (12, 15). GR activation triggers a cascade of events: cation efflux, release of chelated calcium–dipicolinate (Ca-DPA), cortex hydrolysis, and core rehydration, ultimately completing germination (16, 12, 13). In *B. subtilis*, three GRs have been characterized: GerA responds to L-alanine (L-Ala), while GerB and GerK together are activated by a mixture of L-asparagine, D-glucose, D-fructose, and K⁺ (AGFK) (16). Downstream of GR signaling, ion efflux promotes the opening of SpoVA channels, which mediate the bulk release of Ca-DPA (12, 13, 17). In addition to nutrients, certain non-nutrient agents can trigger germination by bypassing specific steps: exogenous Ca-DPA directly activates the cortex-lytic enzyme CwlJ (18), while the surfactant dodecylamine opens SpoVA channels independently of GRs (19, 20).

Notably, high pressure (HP) generated hydrostatically can also trigger spore germination, with the mechanism depending on the pressure magnitude (10, 21). Moderate pressures (50– 300 MPa) are thought to activate GRs (22, 23, 21), whereas higher pressures (>400 MPa) directly open SpoVA channels (22, 23, 21). Despite this phenomenological understanding, the physical basis by which pressure activates GRs or SpoVA channels remains unknown. Among GRs, GerA exhibits the strongest response to moderate HP (24). Intriguingly, although L-Ala and HP both activate GerA, several lines of evidence suggest distinct activation mechanisms (25). For example, heat activation improves L-Ala-triggered germination but does not enhance the HP response (26). Likewise, spores produced at higher sporulation temperatures germinate more efficiently under HP but less efficiently with L-Ala (24). Chemical inhibitors of L-Ala-induced germination, such as ethanol and amiloride, do not suppress HP-induced germination (24). These differences imply that HP activates GerA through a pathway that does not depend on canonical ligand binding.

The GerA complex in *B. subtilis* spores consists of three subunits: GerAA, GerAB, and GerAC (15, 27). Recent structural and functional studies have established that GerA forms a pentameric or hexameric ion-channel complex, in which L-Ala binding to GerAB induces conformational changes that open the pore formed by GerAA (17, 28–30). Building on this framework, we set out to determine how HP activates GerA. Here, we first identify specific residues within the GerA ion-channel region that are essential for HP-induced germination, while showing that classical ligand-binding and intramembrane signaling residues are dispensable. Based on these findings, we propose a stretch-to-open (STO) model in which HP compresses the IM phospholipids, generating lateral membrane tension that mechanically dilates the GerA channel. We test this model through multiple, convergent approaches including *in situ* membrane-tension measurements, computational modeling and calculations, assays of IM permeability under HP, and biophysical modulation of membrane fluidity. Together, these experiments provide convergent evidence that HP activates GerA by compressing the IM and promoting opening of its ion channel, revealing a previously unrecognized mechanosensory mechanism in bacterial spore germination.

## Results

### GerA is the principal target through which high pressure triggers spore germination in *Bacillus* species

A previous study demonstrated that the germinant receptor (GR) GerA and its homologs GerB and GerK play important roles in HP-induced germination at 50–300 MPa, with GerA being the most responsive (24). Consistent with this finding, we observed a similar hierarchy using mutant strains (Δ*gerB* Δ*gerK*, Δ*gerA* Δ*gerK* and Δ*gerA* Δ*gerB*) in the *B. subtilis* PY79 (WT) background (Figure S1A). To precisely define the HP (200 MPa/30°C)-responsive GR in a clean genetic background, we introduced individual *B. subtilis* GRs into the Δ*5* strain, which lacks all functional GRs (Δ*gerBB*, Δ*gerKB*, Δ*yffiT*, Δ*yndE*, and Δ*gerA*) (28), to generate Δ*5 gerA*, Δ*5 gerB* and Δ*5 gerK* (Figure 1D). Notably, only spores expressing GerA (Δ*5 gerA*) germinated efficiently under HP, whereas spores expressing GerB or GerK (Δ*5 gerB*, Δ*5 gerK*), even when overexpressed (Δ*5 P_sspD_-gerB*, Δ*5 P_sspD_-gerK*), showed no response (Figure 1A-1B, 1D). This specificity extended to a GerA homolog from *B. cereus* [Δ5 *gerA (B. c)*], which also conferred efficient HP-induced germination (Figure 1C–1D). Furthermore, deletion of any subunit of GerA (GerAA, GerAB, or GerAC) disrupted GerA complex integrity and eliminated the HP response (Figure S1B-S1C). Together, these results demonstrate that GerA and its functional homolog, but not GerB or GerK, are sufficient to support HP-induced germination in *Bacillus* spores, and that an intact GerA complex is essential for this response.

**Figure 1.**
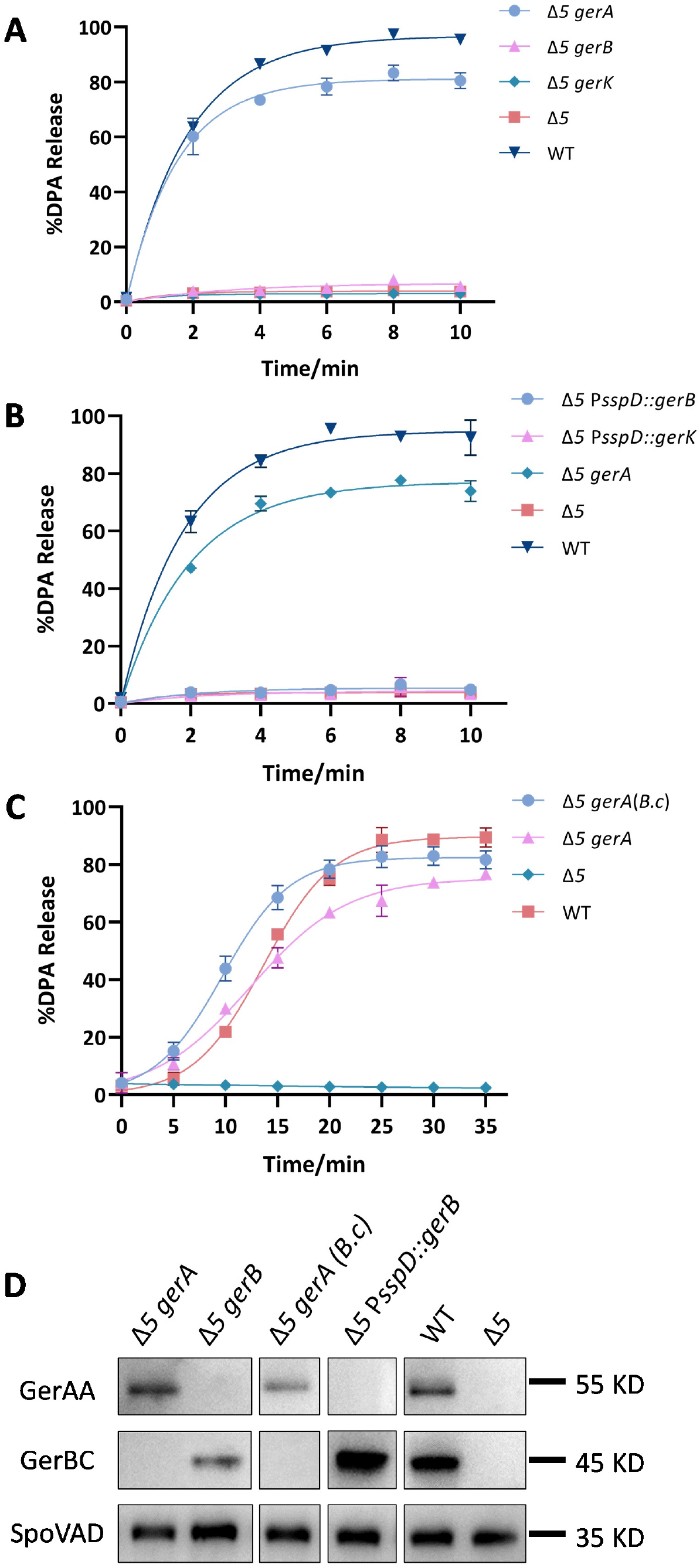
Role of different germinant receptors in HP-induced germination of *Bacillus subtilis* spores. **(A-C)** Spores of TYZ 12 (Δ*5 gerA*), TYZ 13 (Δ*5 gerB*), TYZ 14 (Δ*5 gerK*), TYZ 15 (Δ*5* P*sspD::gerB*), TYZ 16 (Δ*5* P*sspD::gerK*), TYZ 17 [Δ*5 gerA(B. c)*], bLA201 (Δ*5*) and BDR2413 (WT) strains were treated with HP to trigger germination. HP treatments were performed at 200 MPa and 30°C (**A-B**) or 20°C (**C**). DPA was determined by monitoring the relative fluorescence units (RFU) of Tb^3+^-DPA as described in the Materials and Methods, and the percentage of DPA release was calculated as DPA released divided by total DPA content. **(D)** Spores of TYZ 12 (Δ*5 gerA*), TYZ 13 (Δ*5 gerB*), TYZ 15 (Δ*5* P*sspD::gerB*), TYZ 17 [Δ*5 gerA*(*B. c*)], bLA201 (Δ*5*) and BDR2413 (WT) strains were subjected to Western blotting using antibodies against GerAA, GerBC and SpoVAD. A representative result from three independent biological replicates is shown.

### Amino acid residues located in the ion-channel and intracellular regions of GerA are crucial for HP-induced germination

To elucidate the mechanism of HP-induced GerA activation, we leveraged the recent characterization of GerA as an ion channel activated by L-alanine through sequential steps (Figure 2A) (17): (1) ligand binding to GerAB (G25, E105) (28, 29); (2) signal transduction via interfacial residues [GerAA (A386, A313), GerAB (F259)] (28); (3) channel opening through pore-lining residues [GerAA (V362, Q354, Q366)] (17); and (4) downstream signaling via intracellular N-terminal residues (R151, N146, D231) (29, 31). We summarized all these key amino acid residues important for L-Ala-induced activation (Table S1) and examined their roles in HP-induced germination.

**Figure 2.**
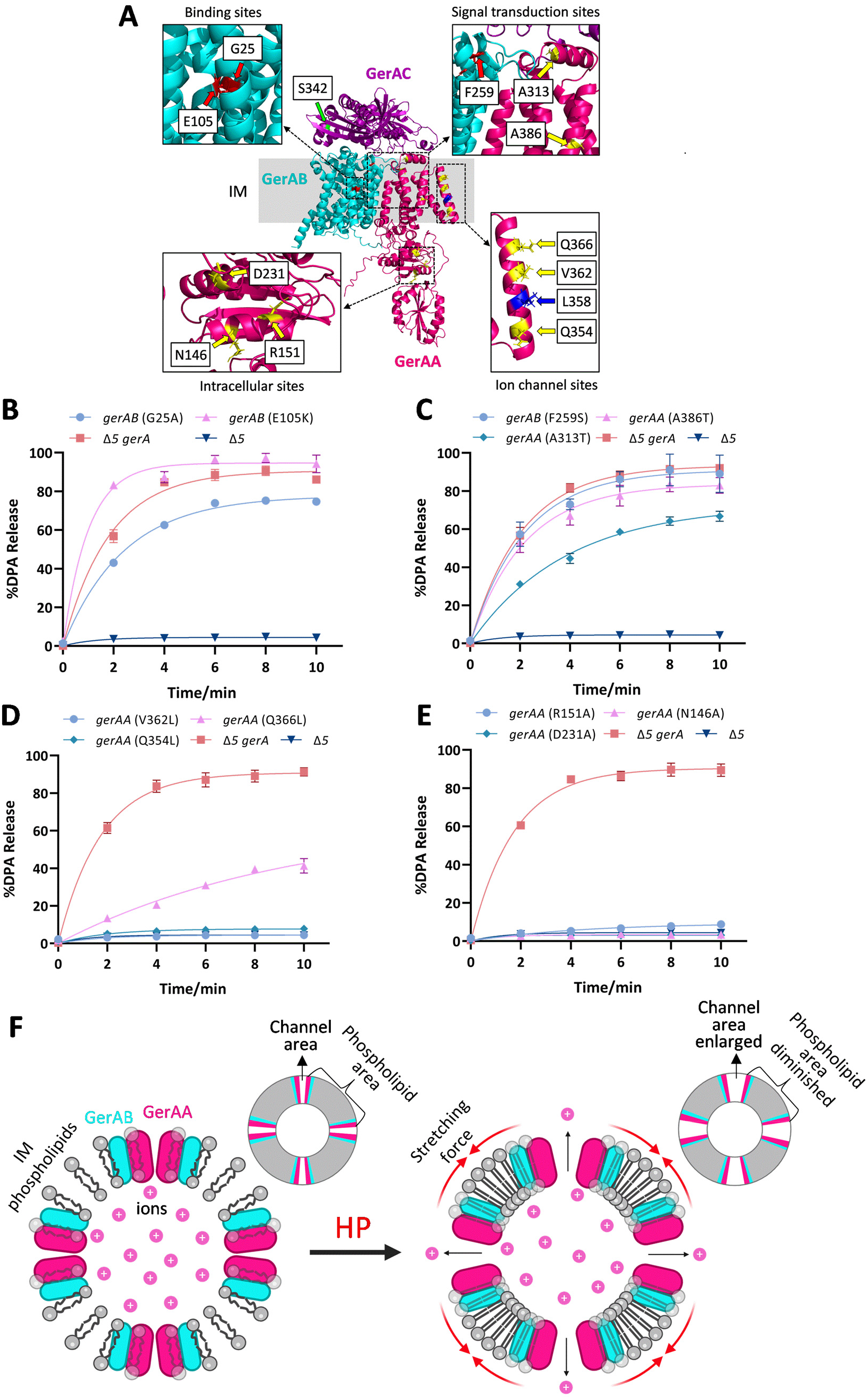
Effects of point mutations in GerA on HP-induced germination. **(A)** Predicted structure of the trimer assembled by GerAA (red), GerAB (cyan), and GerAC (purple). The mutation sites compromising L-Ala-induced germination are categorized into the following categories based on their location: binding sites [gerAB (G25, E105)], signal transduction sites [*gerAB* (F259), *gerAA* (A313, A386)], intracellular sites [*gerAA* (N146, D231, R151)], ion channel sites [*gerAA* (Q366, V362, Q354)], and extracellular sites [*gerAC* (S342)]. **(B-E)** Spores of bLA286 [Δ*5 gerAA/AB(G25A)/AC*], TYZB2 [Δ*5 gerAA/AB(E105K)/AC*], TYZB1[Δ*5 gerAA/AB(F259S)/AC*], TYZA7 [Δ*5 gerAA(A386T)/AB/AC*], TYZA8 [Δ*5 gerAA(A313T)/AB/AC*], TYZA9 [Δ*5 gerAA(V362L)/AB/AC*], TYZA11 [Δ*5 gerAA(Q366L)/AB/AC*], TYZA10 [Δ*5 gerAA(Q354L)/AB/AC*], TYZA2 [Δ*5 gerAA(R151A)/AB/AC*], TYZA4 [Δ*5 gerAA(N146A)/AB/AC*], TYZA5 [Δ*5 gerAA(D231A)/AB/AC*], bLA201 (Δ*5*) and bLA219 (Δ*5 gerAA/AB/AC*) strains were treated with HP at 200 MPa and 30°C to trigger germination. DPA was determined by monitoring the relative fluorescence units (RFU) of Tb^3+^-DPA as described in the Materials and Methods, and the percentage of DPA release was calculated as DPA released divided by total DPA content. **(F)** A mechanistic stretch-to-open (STO) model for HP activation of GerA: High pressure compresses the IM phospholipids more extensively than the core, creating a lateral stretching force across the IM that promotes opening of the GerA ion channel. Each experiment was performed with at least three independent biological replicates.

For this purpose, we constructed point-mutation strains in the Δ*5 gerA* background to alter the key residues described above. In particular, the S342P mutation in GerAC resulted in the loss of GerA complex integrity (Figure S2A-S2B) and consequently caused germination defects with both L-Ala (Figure S2C) and HP (Figure S2D-S2E) (28). Except for *gerAC* (S342P), other point mutations in GerAA and GerAB did not affect GerA complex assembly (Figure S3A, Figure S4A, Figure S5A and Figure S6A), but all mutant spores exhibited severe germination defects when induced by L-Ala (Figure S3B, Figure S4B, Figure S5B and Figure S6B), confirming the expected phenotypic effects of these mutations. When these mutant spores were subjected to HP, mutation or competitive occupation of L-Ala-binding residues [GerAB (G25A, E105K)] (Figure 2B, Figure S3C, Figure S3D-S3E) and mutation of signal-transduction residues [GerAA (A386T, A313T), GerAB (F259S)] (Figure 2C, Figure S4C) caused little or no reduction in HP-induced germination. In contrast, mutation of channel residues [GerAA (V362L, Q354L, and Q366L)] (Figure 2D, Figure S5C) and intracellular residues [GerAA (R151A, N146A, and D231A)] (Figure 2E, Figure S6C) greatly impaired HP-induced germination efficiency. Notably, mutation of residues [GerAA (V362L, Q354L)] (Figure 2A) close to the narrowest site in the channel [GerAA (L358)] (Figure 2A, Dataset S1) exerted stronger inhibitory effects than mutation of the more distal residue [GerAA (Q366L)] (Figure 2D, Figure S5C). These data reveal a distinct genetic requirement: the tested ligand-binding and signal-transduction residues are largely dispensable, whereas residues constituting the ion channel and its intracellular coupling domain are critical for HP-induced GerA activation.

### In situ membrane-tension measurements support a stretch-to-open (STO) model in which HP compresses the spore inner membrane (IM) and promotes GerA opening

The observation that channel-lining residues, but not those embedded within the IM lipid bilayer, are essential for HP-induced germination suggests that HP opens the GerA channel directly, without requiring signal transduction via GerAB. To understand how HP opens GerA, we examined the structural response of the spore to HP. Notably, while the spore core is dense and relatively incompressible (3, 32), the phospholipid-based IM is substantially more compressible (3, 33, 34). Under HP, the IM therefore undergoes greater volumetric contraction than the core it tightly surrounds (for detailed calculations, see Figure S10 and Discussion). This differential compressibility is predicted to generate a lateral stretching force within the membrane plane, analogous to tightening a drumhead. We propose that this membrane tension acts directly on the channel-forming region of GerA to promote its opening. Based on this reasoning, we propose a mechanistic model for HP activation of GerA (Figure 2F): HP compresses the IM phospholipids, creating lateral membrane tension that mechanically stretches and opens the GerA ion channel. This model, referred to as the <u>s</u>tretch-to-<u>o</u>pen (STO) model, is consistent with our mutagenesis data: under HP, the phospholipid bilayer is compressed as a continuum, generating lateral tension. Residues embedded within this lipid environment experience this uniform mechanical force; therefore, mutations at these sites are not expected to substantially alter the overall tension exerted on the channel protein. In contrast, residues in the channel region interact with each other to determine the stretching force required for channel opening; consequently, mutation of these residues strongly affects HP-induced channel opening and subsequent germination.

To test the STO model, we first needed to demonstrate that the IM phospholipids are indeed compressed under HP. To this end, we used the fluorescence probe Flipper-TR, which reports membrane tension changes through its fluorescence lifetime (35), to measure membrane tension in dormant spores under HP in *situ*. To exclude possible membrane tension changes induced by germination, we used Δ*5 gerA* [GerAA (D231A)] spores, which are unable to germinate under HP (Figure 2E, Figure S6C). Moreover, spores were decoated to reduce autofluorescence and to allow efficient probe staining (Figure S7E). As shown in Figure 3A–3B and Figure S7, increasing pressure from 0.1 to 300 MPa significantly elevated the fluorescence lifetime of Flipper-TR *in situ*, consistent with HP-induced phospholipid compression and tension generation. This, combined with the critical role of channel-lining residues (Fig. 2D), supports the hypothesis that HP-induced membrane tension could mechanically facilitate the opening of the GerA channel. If true, elevating pressure should rescue the germination defect of ion-channel residue mutant spores [Δ*5 gerAA* (Q366L), (Q354L), (V362L)] under HP. We then examined the germination of these mutant spores under pressures ranging from 100 to 300 MPa. As expected, the germination defects of these mutant spores were partially rescued, with the effect being particularly pronounced for GerAA (Q354L) and GerAA (Q366L) (Figure 3C). Moreover, the amount of K^+^ released by Δ*5 gerAA* (Q366L) also recovered to a level similar to that released by parental spores (Δ*5 gerA*) (Figure S8), further supporting the functional role of the GerA ion channel. In contrast, intracellular-residue mutant spores [GerAA (N146A), GerAA (R151A) and GerAA (D231A)] failed to recover their germination efficiencies at higher pressures (Figure 3C), indicating that these residues are not direct targets of HP. Interestingly, Δ*5 gerAA* (D231A) spores failed to germinate under HP but achieved substantial K^+^ release (Figure S8), suggesting that these residues are not directly involved in ion release, but likely play important roles in downstream signal transduction after ion release.

**Figure 3.**
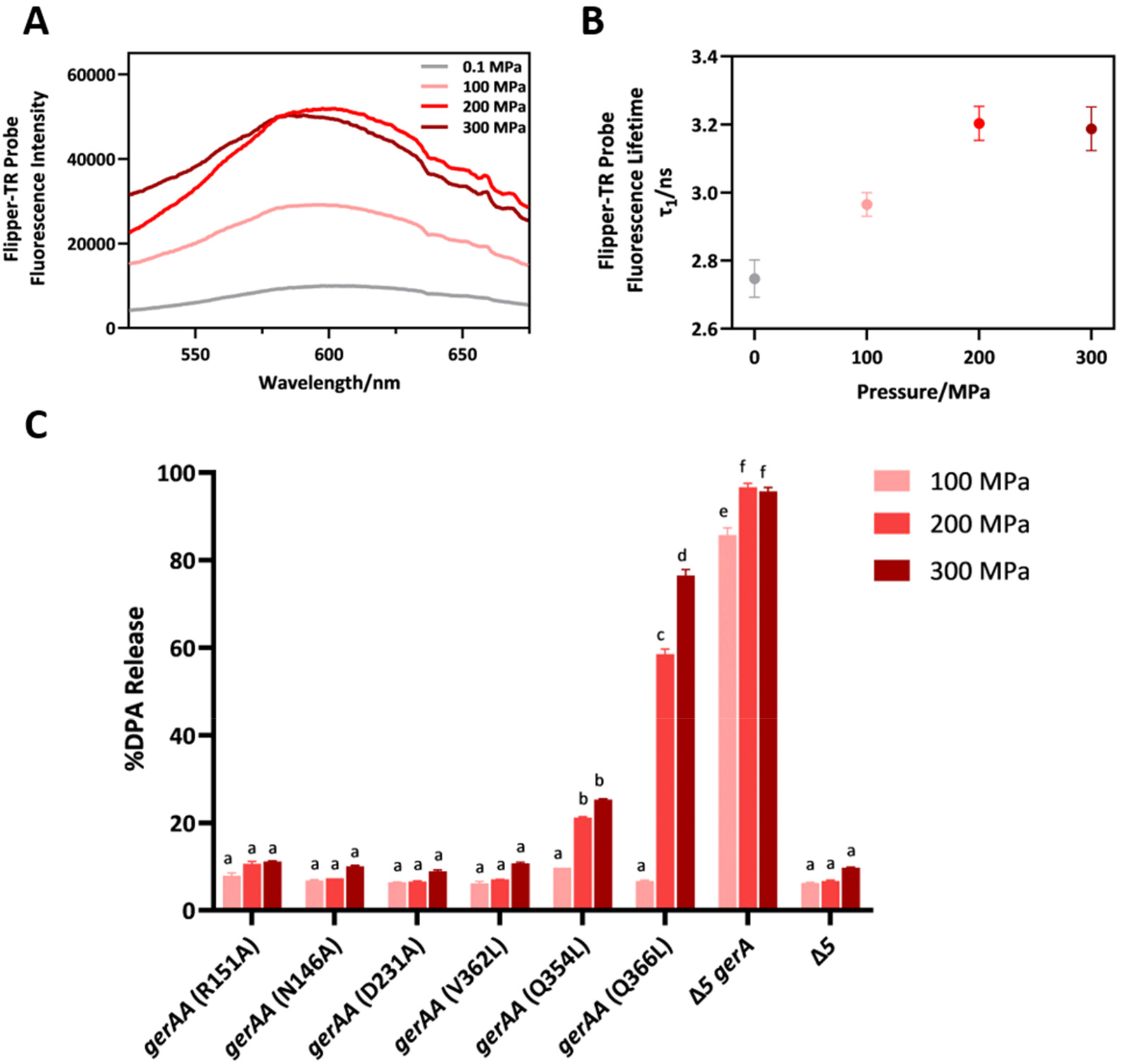
In situ membrane-tension measurements in dormant spores and pressure-dependent germination of GerA point mutants under HP. **(A)** Fluorescence spectra of Flipper-TR probe in the inner membrane of spores TYZA5 [Δ*5 gerAA(D231A)/AB/AC*] at 0.1-300 MPa. The maximum emission peak wavelength of the Flipper-TR probe in the inner membrane under different pressures was within the range of 600/50 nm (detection wavelength of Flipper-TR probe). **(B)** Fluorescence lifetime of Flipper-TR probe in the inner membrane of spores TYZA5 [Δ*5 gerAA(D231A)/AB/AC*] at 0.1-300 MPa. All fluorescence-lifetime fits were performed at the maximum emission wavelength using EasyTau2. **(C)** Germination of spores with mutations in intracellular and ion channel sites under different pressure levels. The spores of TYZA2 [Δ*5 gerAA(R151A)/AB/AC*], TYZA4 [Δ*5 gerAA(N146A)/AB/AC*], TYZA5 [Δ*5 gerAA(D231A)/AB/AC*], TYZA9 [Δ*5 gerAA(V362L)/AB/AC*], TYZA10 [Δ*5 gerAA(Q354L)/AB/AC*], TYZA11 [Δ*5 gerAA(Q366L)/AB/AC*], bLA201 (Δ*5*) and bLA219 (Δ*5 gerAA/AB/AC*) strains were treated at 100, 200, or 300 MPa and 30°C for 20 min. DPA was determined by monitoring the relative fluorescence units (RFU) of Tb^3+^-DPA, as described in the Materials and Methods, and the percentage of DPA release was calculated as DPA released divided by total DPA content. One-way ANOVA with Dunnett’s multiple comparisons test was performed to compare the significant differences. Different letters indicate significant differences between groups (*P* < 0.05). Groups sharing the same letter are not significantly different. Each experiment was performed with at least three independent biological replicates.

Based on the experimental evidence that HP compresses the IM and generates membrane tension (Figure 3A–3B), thereby potentially inducing a stretching effect on GerA (Figure 3C), we performed exploratory molecular dynamics simulations to conceptually examine how lateral tension might affect the GerA channel. Monitoring the pore geometry of GerA at three defined sites (Dataset S1; Q366 at the entrance, L358 at the constriction, and V73 at the exit) suggested a trend of pore enlargement upon the application of a stretching force (Figure 4A–4D, Movie S1). While these computational observations are consistent with a tension-induced GerA opening mechanism, they remain hypothetical and are presented here primarily as a conceptual complement to the experimental data.

**Figure 4.**
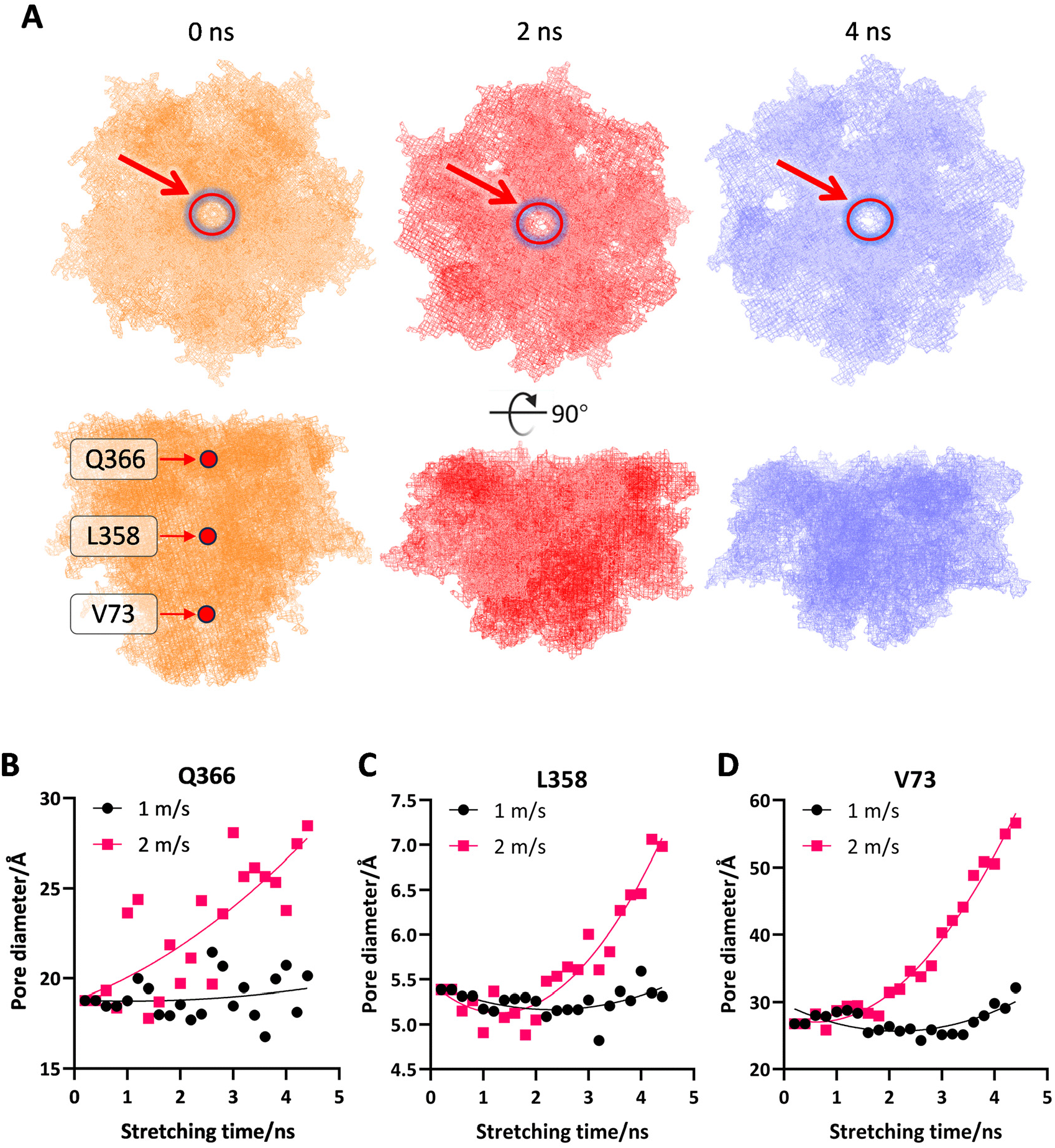
Simulating the opening of GerA under a lateral stretching force. **(A)** After modeling GerA using coarse-grained (CG) methods, lateral stretching of GerA was applied at speeds of 1 m/s and 2 m/s for 0 ns, 2 ns, and 4 ns. The structural changes of GerA are illustrated in both the top (upper) and side (lower) views. The red arrows on the top views indicate the formation of a hole at the center of GerA. Three sites in the GerA ion channel were defined as three local constriction indicators (LCIs) to monitor the opening/closing status of the ion channel: the entrance site (Q366), the narrowest site (L358), and the exit site (V73). **(B)** Changes in the size of the hole at the entrance site (Q366) in the GerA ion channel during stretching processes at speeds of 1 m/s and 2 m/s. **(C)** Changes in the size of the hole at the narrowest site (L358) in the GerA ion channel during stretching processes at speeds of 1 m/s and 2 m/s. **(D)** Changes in the size of the hole at the exit site (V73) in the GerA ion channel during stretching processes at speeds of 1 m/s and 2 m/s. The simulation system was sampled ten times to obtain an average.

### Experimental evidence for HP-induced membrane compression and channel opening further supports the STO model

To obtain additional evidence for the STO model, we examined whether HP increases IM permeability to water-soluble molecules—a direct prediction if membrane compression opens proteinaceous channels. To test this possibility, decoated Δ*5* spores were subjected to HP treatment at 200 MPa in the presence of the water-soluble, membrane-impermeant nucleic acid dye propidium iodide (PI). While untreated spores showed no fluorescence (Figure 5A–5B), HP-treated spores displayed strong red fluorescence in the spore core (Figure 5A–5B), indicating substantial intracellular PI entry. Importantly, these HP-treated spores remained alive, phase bright, and retained all their DPA (Figure 5A, Figure S9A-S9B), arguing against widespread IM damage as the basis for PI entry. Thus, PI likely entered the spore core through the opened protein channels in the IM under HP. We further extended this observation using formaldehyde (CH₂O) (Figure 5C, Figure S9A-S9B), a sporicide that targets core DNA (2). HP significantly enhanced the inactivation of decoated Δ*5* spores by CH₂O, and this effect was further amplified in Δ5 *gerA* [GerAA(D231A)] spores, indicating the opening of the GerA channel (Figure 5C). These results support the conclusion that HP opens GerA as well as other protein channels in the IM. Additionally, the occurrence of K^+^ release from both Δ*5* and Δ*5 gerA* [GerAA (D231A)] spores under 200 MPa (Figure S8) also indicated that ion channels besides GerA and its homologs can be opened by HP. Together, these results support the conclusion that HP increases IM permeability through the opening of GerA and additional ion channels.

**Figure 5.**
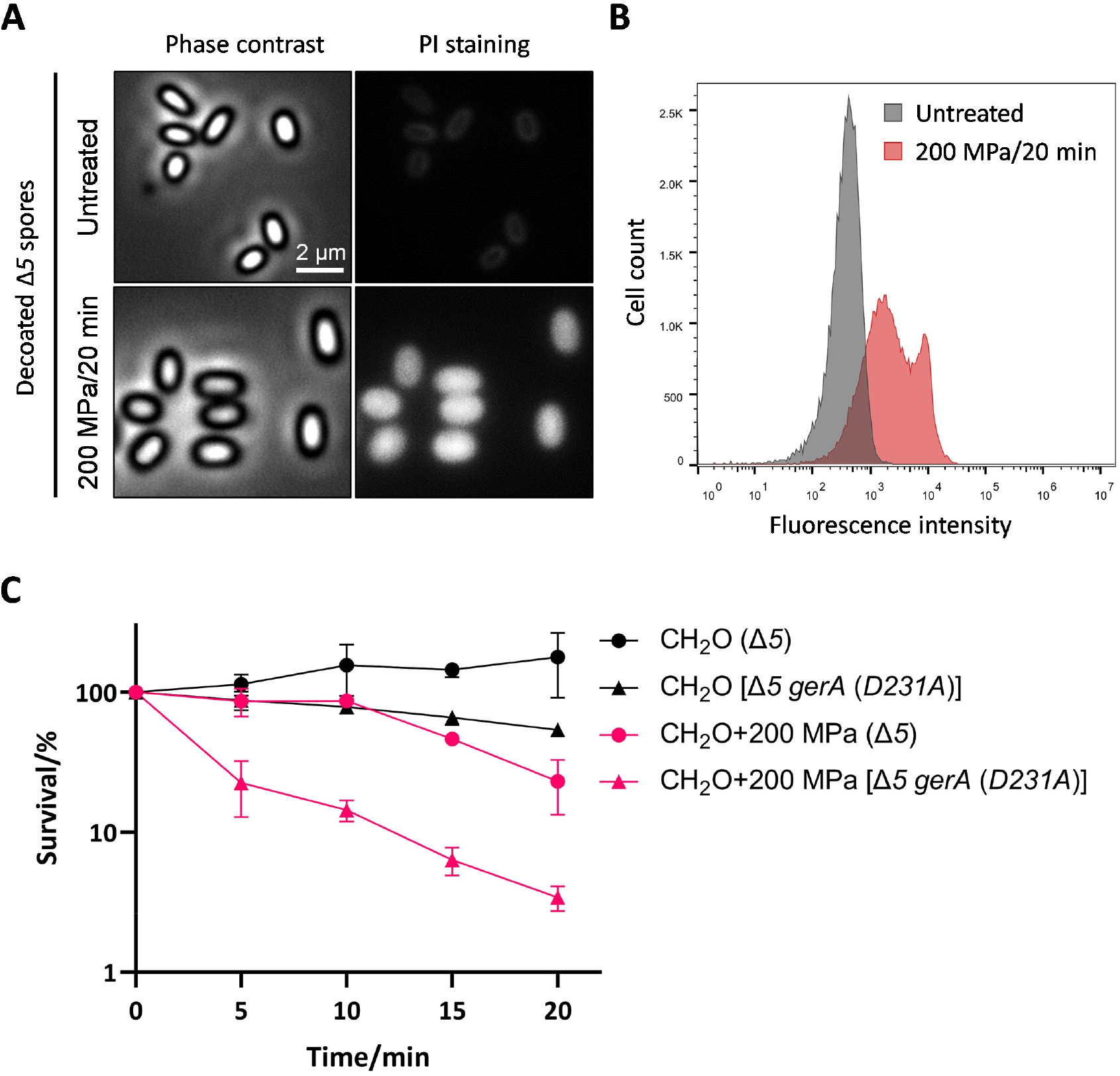
Penetration of water-soluble agents into decoated spores under HP. The bLA201 (Δ*5*) and TYZA5 [Δ*5 gerAA(D231A)/AB/AC*] spores were decoated and tested for their permeability to propidium iodide (PI) and formaldehyde (CH_2_O) under HP at 200 MPa and 30°C, as described in the Materials and Methods. **(A)** Representative phase-contrast and corresponding fluorescence images of the decoated bLA201 (Δ*5*) spores stained with PI. Scale bar, 2 μm. **(B)** Distributions of the mean per-cell fluorescence of the untreated decoated Δ*5* spores (gray) and 200 MPa/20 min-treated decoated Δ*5* (red) cells stained with PI. **(C)** Survival of the decoated Δ*5* and Δ*5 gerA* [*gerAA* (*D231A*)] (simplified as Δ*5 gerA* (*D231A*) in the figure) spores treated with 2.5% CH_2_O under 0.1 MPa or 200 MPa at 30°C. Each experiment was performed with at least three independent biological replicates.

We next tested a central prediction of the STO model: that GerA opening results from HP-induced membrane compression. If true, the physical state of the IM should govern its compressibility and thus the efficiency of GerA activation. Specifically, an IM with higher fluidity and more loosely arranged phospholipids possesses lower intrinsic membrane tension, and should therefore require higher pressure to generate sufficient tension for channel opening (Figure 6E). Exploiting this theory, we increased IM fluidity by heat activation or chemical decoating of Δ5 *gerA* spores (26, 36), as indicated by a reduction in Laurdan generalized polarization (GP) values (Figure 6A–6B) (37). Direct tension measurements using Flipper-TR corroborated that heat-activated spores exhibited reduced intrinsic IM tension (Figure 6B, Figure S9C-9E). As predicted by the model, these heat-activated or decoated spores with elevated IM fluidity demonstrated markedly delayed germination kinetics under 200 MPa HP treatment compared to untreated spores (Figure 6C–6D). These findings support the IM as a key mechanical transducer of HP and identify phospholipid compression as the critical event driving GerA channel activation (Figure 6E). In summary, our integrated experimental approach combining permeability assays, biophysical tension measurements, and membrane-state modulation provides strong, consistent evidence that HP compresses the IM to generate lateral tension that promotes opening of the GerA ion channel via the proposed STO mechanism.

**Figure 6.**
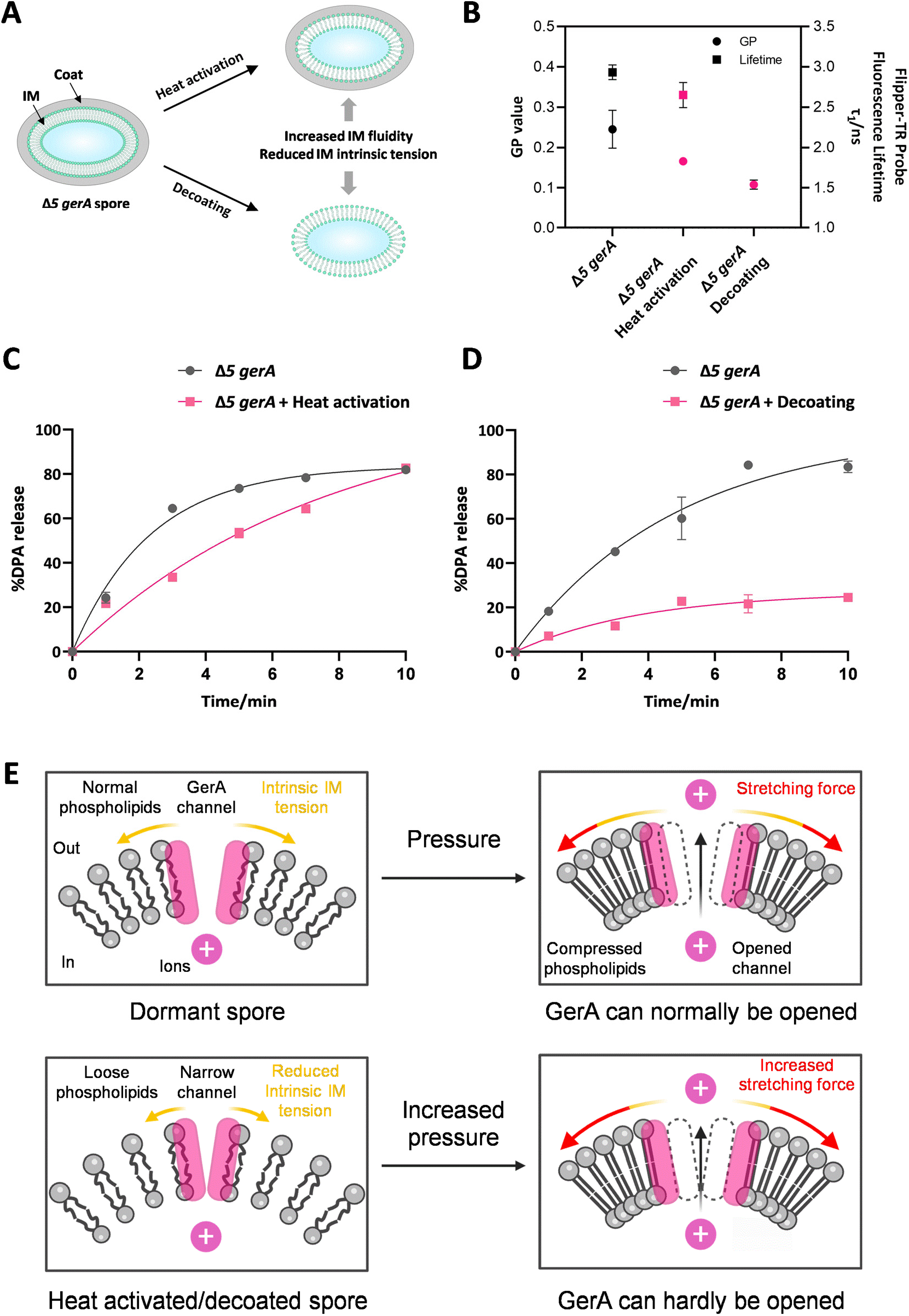
Effect of heat activation or decoating on HP-induced germination of spores. The bLA219 (Δ*5 gerAA/AB/AC*) (simplified as Δ*5 gerA* in the figures) spores were heat activated (75°C/60 min) or chemically decoated (see Materials and Methods) to modify the inner membrane (IM) fluidity and intrinsic tension **(A)**. The spore IM fluidity and tension were assessed by measuring the generalized polarization (GP) of the fluorescent dye Laurdan and the fluorescence lifetime of the Flipper-TR probe incorporated into the spore IM **(B)**, as described in the Materials and Methods. The heat-activated **(C)** or decoated **(D)** Δ*5 gerA* spores were treated with HP at 200 MPa and 30°C to trigger germination. DPA was determined by monitoring the relative fluorescence units (RFU) of Tb^3+^-DPA as described in the Materials and Methods, and the percentage of DPA release was calculated as DPA released divided by total DPA content. Each experiment was performed with at least three independent biological replicates. **(E)** Schematic illustration of the HP-induced ion channel activation in the TYZ12 (Δ*5 gerA*) spores subjected to heat activation or decoating. The pink bars represent the GerA channel, the gray sections represent the phospholipid bilayer, and the pink spheres represent the ions. The yellow arrows and red arrows indicate the IM intrinsic tension and HP-induced stretching force, respectively. The hollow dashed bars indicate the original position of the subunits forming the GerA channel before HP treatment.

## Discussion

The ability of high pressure (HP) to induce bacterial spore germination has been recognized for decades (38, 39). While subsequent studies established that moderate pressures (50–300 MPa) act primarily through germinant receptors (GRs) and very high pressures (>400 MPa) directly open SpoVA channels (23, 24), the precise biophysical mechanism by which HP activates these membrane proteins has remained elusive. In this study, we provide convergent evidence supporting a stretch-to-open (STO) model, whereby HP compresses the inner membrane (IM) to generate lateral tension that promotes dilation of the GerA ion channel, thereby initiating germination (Figure 7). This model directly links an applied physical force to the conformational gating of a key germination receptor of bacterial spores.

**Figure 7.**
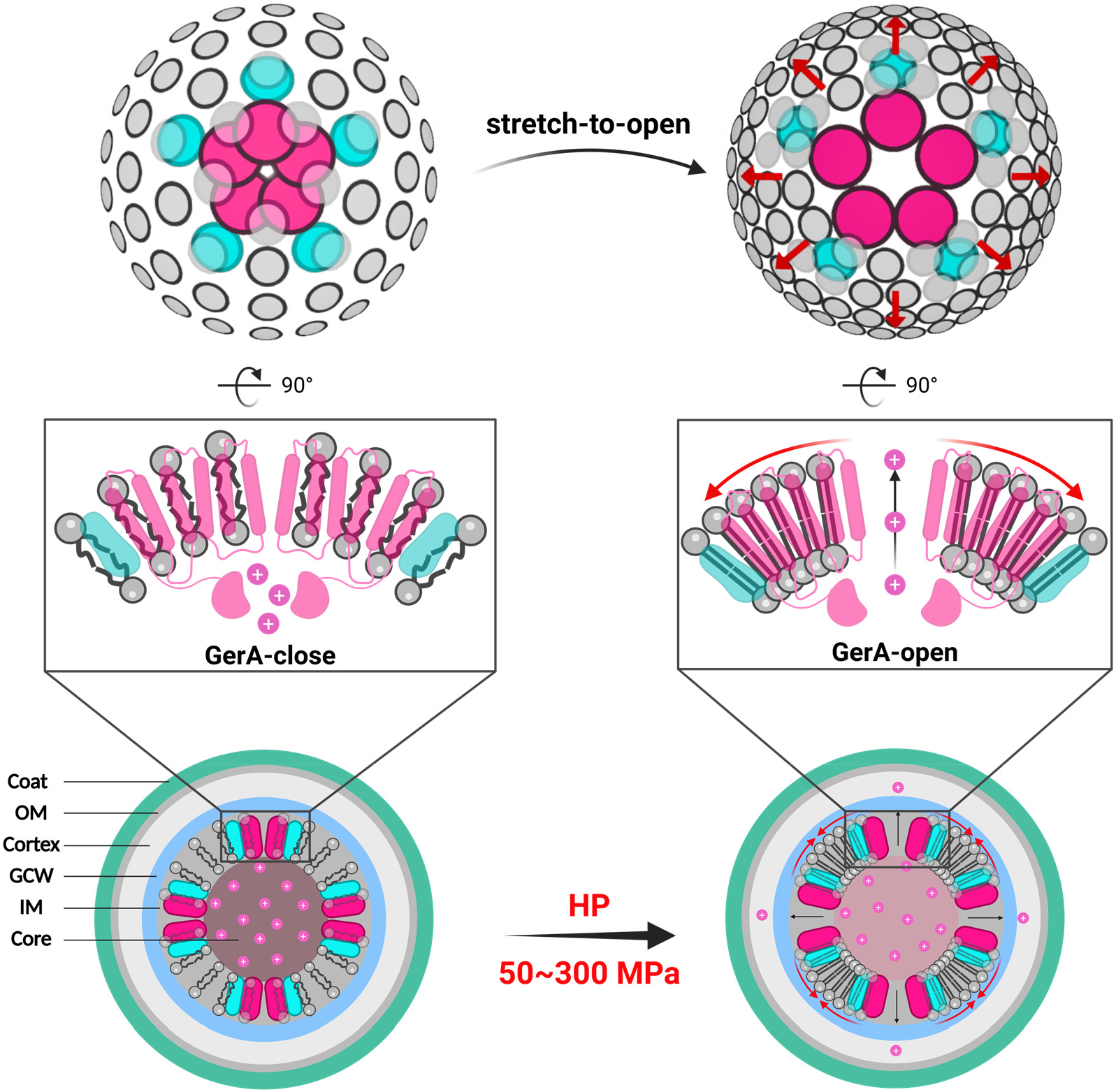
Stretch-to-open model for HP-induced GerA activation. Schematic illustration of the activation of the ion channel GerA under moderate HP (50–300 MPa) through the compression of the inner membrane (IM) phospholipids. Specifically, HP compresses the IM phospholipids, generating a lateral stretching force that promotes opening of the GerA ion channel. The pink sections represent the GerAA subunit, the teal sections represent the GerAB subunit, the gray sections represent the phospholipid bilayer, and the pink spheres represent the ions. The red arrows on the top view indicate the stretching direction in which GerA is displaced as phospholipids are compressed. OM, outer membrane; GCW, germ cell wall; IM, inner membrane.

Regarding this new model, a central question is how phospholipid compression translates into a membrane-plane stretching force. HP reduces intermolecular distances, and the degree of compression depends on a material’s compressibility (40). For bacterial spores, the IM is constructed from phospholipid molecules that assemble via noncovalent bonds into a phospholipid bilayer interwoven with proteins (3), exhibiting considerable structural flexibility. In contrast, the spore core is packed with dense compounds like Ca-DPA and metal cations, making it far less compressible (3, 32). We therefore hypothesized that the IM would compress more readily than the core under HP, and that this differential compressibility would generate lateral tensile stress in the membrane. To examine this hypothesis, we constructed a simplified physical model of the spore, featuring a core (approximated as water) surrounded by a DMPC (1,2-Dimyristoyl-sn-glycero-3-phosphocholine) bilayer containing GerA and SpoVA (17, 41) (Figure S10A). Volume calculations across a pressure range of 0.1–600 MPa predicted a distinct crossover (Figure S10B): at moderate pressures (≤300 MPa), the IM volume decreased faster than the core volume, whereas at very high pressures (≥400 MPa), the core volume decreased faster. This differential drives the mechanical response. In the moderate-pressure regime, the faster-contracting IM experiences lateral compression, generating a stretching force that dilates and opens the GerA channel (Figure S10C). Concurrently, this expansion tightens the D-subunit plug in the SpoVA(C-D-Eb) CaDPA channel, preventing its activation (Figure S10C). Conversely, in the high-pressure regime, the faster-contracting core radially compresses the IM. This inward compression squeezes the GerA channel closed while displacing the SpoVAD plug, thereby opening the SpoVA (C-D-Eb) channel (Figure S10C). We note that modeling the core as water, a more compressible medium than the actual mineral-rich core, provides a conservative estimate; the true core is less compressible (blue dashed line, Figure S10B), which would raise the absolute pressure thresholds but preserve the fundamental crossover logic. Thus, these calculations provide a quantitative framework for the STO model that may help explain the two established pressure-response pathways.

The STO model posits that the physical state of the IM and the conformation of the channel region are primary determinants of HP-induced germination. This framework elegantly explains long-standing disparities between L-Ala- and HP-induced germination (10, 25): (1) D-alanine competitively inhibits L-Ala-induced germination but does not affect HP-induced germination (Figure S3D-S3E) (42), because HP acts via membrane tension rather than the specific ligand-binding pocket; (2) Heat activation improves L-Ala-induced germination but not HP-induced germination (26). This occurs because the heat treatment temporarily increases the IM fluidity, causing a looser arrangement of the phospholipids (34). This enhances the frequency with which L-Ala binding pockets are exposed (29). However, increased IM fluidity also increases the pressure required to compress phospholipids, thereby producing the opposite effect on HP-induced germination (Figure 6A, 6C, 6E). This also explains why spores formed at higher temperatures (with less fluid IMs) germinate better under HP but worse with L-Ala (24); (3) Ethanol inhibits L-Ala-induced germination but not HP-induced germination. Ethanol itself compresses and tenses the IM (43, 44); this reduces the exposure of the binding pockets of GerAB and thus inhibits L-Ala-induced germination. Conversely, this ethanol-induced compression of the IM lowers the pressure required to induce phospholipid compression in HP-induced germination, producing a positive effect; (4) Spores activated by HP (150 MPa/30 s) require continuous external pressure to maintain the activation state of GerA (45). This likely occurs because the external pressure is needed to maintain the compression of the IM phospholipids and subsequent channel opening.

An unexpected finding in this study is that GerB and GerK, which AlphaFold predicts to be structurally similar to the GerA ion channel (46), are unable to independently respond to HP (Figure 1A–1B). This is consistent with our recent observation that the SpoVAF/FigP ion channel, which shares a GerAA-like fold (47), is also HP-insensitive (10); these results indicate that structural similarity to GerA is not sufficient for pressure sensing. Interestingly, it has been reported that GerB and GerK are capable of responding to HP in the presence of accessory proteins YfkT and YndE (Figure S1A) (24), suggesting that functional mechanosensitivity in this receptor family may require specific cofactors or molecular contexts that remain to be defined.

Collectively, our evidence positions GerA as a mechanosensitive ion channel gated by membrane tension. Mechanosensitive channels are ubiquitous in bacteria and vital for responding to osmotic and mechanical stress (48–50). Notably, the *E. coli* MscK channel, which is also activated by lateral membrane tension and shares structural features with GerA, is essential for survival under high-K⁺ conditions (49). Interestingly, GerA appears less sensitive than many canonical mechanosensors. The evolutionary basis of this property remains to be established. Sporulation is an ancient adaptation, likely originating in marine environments characterized by consistently elevated hydrostatic pressure (51–53). GerA may therefore have evolved as a specialized, moderately sensitive hydrostatic pressure sensor. In contrast, other mechanosensors may respond at lower tension thresholds, as exemplified by MscK’s response to mild osmotic gradients (54, 55).

## Materials and Methods

### Strains and general methods

The *B. subtilis* strains used in this study are derivatives of the wild-type strains PY79 and BDR2413, as detailed in Table S2. The construction of plasmids and the specific primers used are described in Tables S3 and S4, respectively. All standard procedures for handling *B. subtilis* followed the methods outlined by Harwood and Cutting (56). For sporulation, cells were incubated at 37°C in Schaeffer’s liquid medium (Difco Sporulation Medium, DSM) (56). Mature spores were collected using the purification method established previously (57). Before germination assays with nutrient germinants, purified spores were heat-activated at 75°C for 30 minutes. Nutrient-germination was then induced by adding either 10 mM L-alanine or a mixture containing 2.5 mM L-asparagine, 5 mg/mL D-glucose, 5 mg/mL D-fructose, and 50 mM KCl, with incubation continuing at 37°C in 25 mM K-Hepes buffer (pH 7.4).

### Spore purification

The purification of mature *B. subtilis* spores was carried out using the method described previously (57). First, a 22-hour culture grown in Difco Sporulation Medium (DSM) was centrifuged, and the cells were washed three times with double-distilled water (ddH_2_O). The washed cells were then stored at 4°C under constant agitation. Each day, the suspension was washed and resuspended in fresh ddH_2_O. After seven days, spores were collected by centrifugation to obtain a pellet. The pellet was then resuspended in a 20% Nycodenz solution, using 400 μL of the solution for every 10 mL of the original DSM culture, and left on ice for 30 minutes. From this mixture, 200 μL aliquots were layered onto 900 μL of a 50% Nycodenz solution and subjected to gradient fractionation by centrifugation at 20,000 *g* at 4°C for 30 minutes. The resulting pellet was washed at least five times with ddH_2_O. The purity of the spores was assessed using phase-contrast microscopy, and only those spores with greater than 99% purity were used in subsequent experiments. If the required purity was not achieved, the purification process was repeated.

### High-pressure-induced spore germination

Aliquots (1.5 mL) of spores suspended in PBS (OD_600_ = 1) were sealed in sterile, flexible plastic bags. High-pressure (HP) treatments were conducted using a FPG7100:9/2C high-pressure isostatic system (Stansted Fluid Power Ltd., Harlow, Essex, UK) equipped with a 2.5-liter pressure vessel. Water served as the pressure-transmitting fluid and temperature regulator. For each treatment, spore samples were loaded into the vessel and subjected to pressurization, pressure holding, and depressurization. The pressurization and depressurization durations were fixed at 2 min and 2 s, respectively, with the pressure-holding time defined as the effective HP treatment duration. Spore suspensions were processed at designated pressures and 30°C for varying holding time intervals (2, 4, 6, 8, and 10 min). Immediately after treatment, samples were chilled on ice and centrifuged (12,000 × g, 10 min, 4°C) within 1 h. For dipicolinic acid (DPA) quantification, 198 μL of supernatant was mixed with 2 μL of 5 mM terbium chloride (TbCl_3_) in 96-well plates. The fluorescence intensity of the Tb³⁺-DPA complex was measured (excitation/emission = 270/545 nm) using a Spark 10M microplate reader (Tecan, Switzerland). To assess spore germination, the centrifuged pellets were examined for phase-dark spores via phase-contrast microscopy (Nikon DS-Qi2 camera with a Plan Apo Lambda 100×/1.45 oil immersion objective; Nikon, Japan).

### DPA measurements

DPA release was detected as described previously with some modifications (58). Briefly, spore germination was induced by L-Ala at 37°C in a 96-well plate. Spores at an OD_600_ of 0.5, 10 mM germinants, 25 mM K-Hepes buffer (pH 7.4), and 50 μM TbCl_3_ were mixed in a total volume of 200 μL. The fluorescence intensity of the Tb³⁺-DPA complex was monitored at Ex/Em = 270/545 nm using a TECAN Spark 10M microplate reader (TECAN, Switzerland). The total DPA content of the spores was measured by boiling spores (OD_600_ of 1) for 20 minutes, then mixing the boiled spores with 50 μM TbCl₃ in a 200 μL solution in a 96-well plate. A DPA standard solution was serially diluted and measured alongside the samples to generate a standard curve. The detection parameters for DPA release were identical to those used for germination assays, and the total DPA content of the spores was calculated based on the standard curve.

### Quantification of potassium-ion release

Potassium-ion release from spores under moderate HP was measured using Agilent 7800 ICP-MS (Agilent Technologies, Inc., USA) by Shiyanjia Lab (Beijing, China). A 2.5 mL aliquot of purified spore suspension (OD_600_ = 2) was sealed in each flexible plastic bag. These bags were subjected to high-pressure treatment at 200 MPa and 30°C for 20 minutes. For quantification of total potassium ions in spores, the spore suspensions were transferred to FastPrep tubes containing Lysing Matrix B beads and lysed with a FastPrep-24 (MP Biomedicals, LLC, USA) at a speed setting of 6.5 for 3 × 60 s, with tubes kept on ice between runs. The supernatants from the pressure-treated and lysed spore suspensions were collected by centrifugation at 15,000 *g* for 10 minutes at 4°C and stored on ice. The supernatants were then filtered through 0.2 μm syringe filters to remove particulates. 1 mL of the supernatant was pipetted into a polytetrafluoroethylene beaker containing 10 mL of 68% nitric acid and heated on a 200°C electric heating plate until organic matter was fully digested. When 1 mL of digestion solution remained, it was diluted with ddH_2_O to a final volume of 10 mL and then analyzed by ICP-MS. The concentration of potassium was determined based on a standard curve generated using standard substances (TMRM^®^ Co., Ltd., Beijing, China). Potassium release was expressed relative to the total potassium content determined after lysis.

### In situ measurement of membrane tension under high pressure

Membrane tension of the spore inner membrane was characterized by the fluorescence lifetime of a Flipper-TR probe under high pressure *in situ*. Decoated spores (OD_600_=100) were stained with 25 μM Flipper-TR probe (SpiroChrome, Swiss) at 37°C for 30 min. *In situ* high-pressure fluorescence spectra and fluorescence lifetime measurements were performed using a spectroscopy system (Ideaoptics, China) developed by the Center for High Pressure Science and Technology Advanced Research. The system utilized a 405 nm laser as the excitation source. A symmetric diamond anvil cell, equipped with type IIa diamonds polished to a diameter of 400 µm, was employed to generate the high-pressure environment. Steel gaskets were pre-indented to a thickness of approximately 40 µm, and 200 µm holes were drilled into them to serve as sample chambers. Stained spores were placed in the sample chamber together with ruby spheres, thereby applying hydrostatic pressure to the sample. Pressure calibration was performed based on the fluorescence peak of the ruby spheres (59). The fluorescence lifetime was fitted using a double-exponential model with EasyTau2 software, and the longest lifetime with the higher fit amplitude (τ1) was used to report the membrane tension (35).

### Phase-contrast/fluorescence microscopy and flow-cytometry analyses

Phase-contrast and fluorescence microscopy were performed using a Nikon DS-Qi2 microscope equipped with a Nikon Plan Apo Lambda 100x/1.45 Oil Microscope Objective. For propidium iodide (PI) staining, the decoated Δ*5* spores at an OD_600_ of 1 were stained with 10 μg/mL PI for 20 min in the dark. Then, 50 μL of spore suspension was centrifuged, and the pellet was resuspended in 5-10 μL of 1x PBS and imaged. For unstained controls, 50 μL of spore suspension at an OD600 of 1 was centrifuged, and the pellet was resuspended in 5-10 μL of 1 x PBS and imaged. Image analysis was conducted using ImageJ2 software. The PI-stained spores were also analyzed using Accuri C6 flow cytometry (Becton, Dickinson and Company, USA). To ensure the accurate collection of spores, the forward-scatter threshold was set to 5000. The sample collection flow rate was 14 µL/min and the fluidic-core diameter was 10 µm. 50,000 events were collected from each sample and analyzed by FlowJo V10.

### Spore inactivation by sporicidal agents

Each 2.5 mL aliquot of decoated spore suspension (OD_600_ = 2) was sealed in a flexible plastic bag with 2.5% formaldehyde (60). These bags were subjected to HP treatment at 200 MPa and 30°C for 0, 5, 10, 15, or 20 min. The treated spore suspensions were then diluted with 50 mmol L⁻¹ PBS buffer (pH 7.4) and spread onto LB plates, followed by incubation at 37°C. Colonies were counted after no additional colonies appeared. The data were plotted as the survival percentage over time.

### Spore decoating

Dormant spores at an OD_600_ of 50 were decoated in decoating buffer [50 mM Tris-HCl (pH 7.4), 8 M Urea, 1% SDS and 50 mM DTT] at 37°C for 1 h with rotation (61). The spore suspension was centrifuged at 20,000 × *g* for 5 min, and the supernatant was removed before the pellet was washed with SDDW at least five times. A 500-μL aliquot of decoated spores at an OD600 of 1 was then incubated at 37°C for 20 min in 0.85% NaCl containing 25 mg/mL lysozyme. Successful decoating was confirmed by measuring the decrease in OD600 during lysozyme-induced germination and by phase-contrast microscopy. The decoated spores were then suspended in ddH_2_O and stored at 4°C.

### Measurement of spore inner-membrane fluidity

The inner membrane (IM) fluidity of spores was quantified by measuring the generalized polarization (GP) of the fluorescent dye Laurdan incorporated into the IM as described previously (62, 37). Briefly, Δ*5 gerA* strain was cultured in Luria-Bertani broth overnight at 30°C, and 200 μL of the culture was spread onto 100 μM Laurdan-containing DSM agar and incubated at 37°C in the dark for 2-3 days. Mature spores were harvested using a cell scraper and purified as described above. Spore suspensions (OD_600_ = 0.5) were aliquoted (200 μL/well) into a 96-well plate. Spore IM fluidity was quantified via the GP value using the following equation:

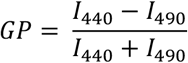

where I_440_ and I_490_ were the fluorescence emission intensities at 440 nm and 490 nm, respectively, measured following an excitation at 360 nm using a TECAN Spark 10M microplate reader (TECAN, Switzerland).

### Construction of point-mutant strains

To replace the original allelic fragment of the target gene, primers were selected to amplify GerAA, GerAB, and GerAC, each with 500-bp upstream and downstream flanking sequences. Both the amplified target fragments and the vector pDG364 were digested with the restriction enzyme BamHI (NEB). The target fragments were then ligated into the plasmid pDG364 using Quick Ligation (NEB). Complementary primers were designed so that the codon of the mutant amino acid was positioned in the middle of the primer sequence, allowing for the replacement of the original codon with the codon of the target amino acid. Then, the full-length plasmid containing the desired mutation was obtained via inverse PCR amplification. Subsequently, the DNA template was digested with DpnI (NEB) at 37 °C for 2 h to ensure complete removal of the parental plasmid. Successful introduction of the mutation was confirmed by DNA sequencing. The full-length target fragment containing the mutation site was then amplified by PCR and assembled with an antibiotic-resistance cassette and the downstream fragment of the original gene using Gibson Assembly. The assembled fragments were then transformed into *B. subtilis* strain bLA219 (Δ*5 gerA*). Colonies growing on the corresponding selective antibiotic plates were selected for genomic DNA extraction, followed by PCR amplification and sequencing to confirm the presence of the point mutation.

### Western blotting

Spores were resuspended in 1x PBS supplemented with 0.05% SDS and Halt Protease Inhibitor (Pierce). Spore lysis was achieved using the FastPrep system (MP, 6.5 m/s, 60 seconds, three cycles). Following lysis, the samples were centrifuged at 20,000 *g* for 10 minutes at 4°C, and the resulting supernatant was used for Western blotting. Protein extracts were mixed with Laemmli sample buffer and heated at 100°C for 10 minutes. Proteins were then separated on a 12.5% SDS-PAGE gel and transferred onto a polyvinylidene difluoride (PVDF) membrane (Immobilon-P, Millipore). The membrane was blocked in a solution of 0.1% Tween-20 and 3% BSA in 1x TBS, followed by a 2-hour incubation with primary antibodies: anti-GerAA (1:5,000), anti-GerBA (1:5,000), and anti-SpoVAD (1:10,000) (63). Afterward, the membrane was incubated for 1 hour with an anti-rabbit IgG HRP-linked secondary antibody (1:3,500). Each antibody incubation was performed in a blocking buffer appropriate for the respective protein to ensure optimal binding and minimal non-specific interactions.

### Molecular dynamics simulations

Following the method described by Gao et al.(17), we used AlphaFold-Multimer to model the multimeric structure of GerA, submitting the protein sequence with default parameters (46). According to the method previously described (64, 65), we employed coarse-grained (CG) models where small groups of atoms are represented as single particles. This approach allowed us to extend the timescale of the membrane-stretching simulations to capture the entire process. First, we constructed a membrane model containing 1152 phosphatidylethanolamine lipids. Then, we placed the GerA multimer centrally within this membrane inside a simulation box with dimensions of 28 × 28 × 28 nm (perpendicular to the Z-axis). The box also contained approximately 44,869 coarse-grained water molecules. After energy minimization, we performed a 1 μs equilibrium simulation of the initial system using an isothermal-isobaric (NPT) ensemble with periodic boundary conditions and a timestep of 20 fs. We then employed the Berendsen thermostat method (66) to maintain the system at 300 K with a coupling time of 1.0 ps. The same algorithm was also used to control the system pressure, set at a reference pressure of 1 bar, with a coupling constant of 4 ps and a compressibility of 3 × 10^-5^ bar^-1^. The stretching process followed the method outlined by Koshiyama and Wada (67), utilizing non-equilibrium stretching simulations. All simulations were conducted using the ‘deform’ option in the GROMACS 5.1.2 software. This method involves scaling the box lengths *l*_i_ and all atomic coordinates *r* proportionally with the timestep 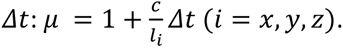. In the simulations presented, the same scaling factor *μ* is used for equal biaxial stretching in the *x* and *y* directions. Here, *c* represents the stretching speed in the *x* or *y* direction of the simulation box, set at 1 m/s (80 mN/m) and 2 m/s (95 mN/m), respectively. During the stretching process, the Parrinello-Rahman barostat maintains the pressure along the *z*-direction at 1 bar, with a coupling constant of 2 ps and a compressibility of 4.5×10^−5^ bar^−1^, allowing the system to contract freely in the *z*-direction. The other parameters of the stretching simulation remain consistent with those used in the equilibrium simulation. Due to the non-steady-state nature of the stretching process, the simulation system is sampled ten times to obtain an average. Visualization was performed using Visual Molecular Dynamics (VMD) 1.9.3 and PyMOL 2.4.0.

### Mathematical calculation of HP-induced volume changes in the spore core and IM

First, we simplified the spore structure to construct a model including the spore core and phospholipid bilayer embedded with GerA and SpoVA using Adobe Illustrator (Adobe, USA). In this model, the material inside the spore core was approximated as pure water, and the phospholipid bilayer was DMPC (1,2-dimyristoyl-sn-glycero-3-phosphocholine), tightly adhering to the spore core. The spore core was approximated as an ellipsoid, with the mathematical expression as follows (Figure S10A).

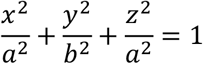

Under HP treatment at 30°C, the relative volume of the spore core as a function of pressure (*V*_core,p_), could be calculated using Eq. 1.

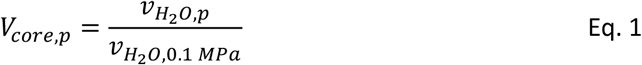

Here, 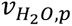 was the specific volume of water at pressure *p*, and 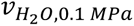 was the specific volume of water at 0.1 MPa. The specific values of 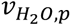 were obtained from Fluid Physical and Chemical Properties Data Resource Platform (http://thermodata.cn/calculation/), and *V*_core,p_ values were calculated by Eq. 1 and presented as the solid blue line in Figure S10B.

To calculate the volume change of the phospholipid bilayer, the thickness of the bilayer was set to be *d*. Since this thickness *d* is much smaller than the size of the spore core (*d* « *a*, *d* « *b*) (68), the volume of the phospholipid bilayer was approximated as the product of the spore core’s surface area and the thickness *d* of the phospholipid bilayer, calculated using Eq. 2.

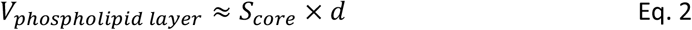

The surface of the spore core was formed by *N* tightly packed phospholipid molecules, with each phospholipid molecule approximated as a cylinder with radius of *r* and height of *d*/2. The area occupied by each phospholipid molecule could be considered as an area element *dA*, which could be approximated as a square with side length of *2r*. Thus, the surface area of the spore core could be calculated using Eq. 3.

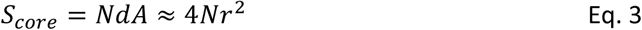

According to previous studies, under HP conditions, the apparent specific volume of a single phospholipid ( *v_phospholipid,p_*) molecule decreased, with lateral compression and vertical stretching of the phospholipid molecules (33, 34). We approximated that the radius of the phospholipid cylinder became *k*_1_ of the original, and the height became *k*_2_ of the original (Figure S10A). The relative volume of a single phospholipid under different pressures (*V_phospholipid,p_*) could be expressed as the ratio of the specific volume of phospholipid under different pressures ( *v_phospholipid,p_*) to the specific volume of phospholipid at 0.1 MPa (*v_phospholipid_*_,0.1_ *_MPa_*), calculated using Eq. 4.

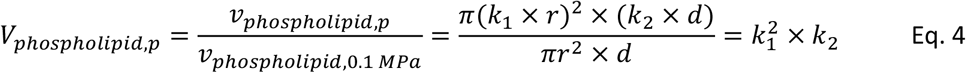

For the phospholipid bilayer, under HP treatment at 30°C, the total number of phospholipid molecules *N* remained unchanged. Since the radius of the approximated cylinder became *k*_1_ of the original, the area element *dA’* occupied by each phospholipid molecule became 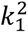 of the original, and the surface area under HP was calculated using Eq. 5.

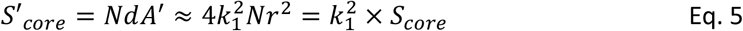

The volume of the compressed phospholipid bilayer was approximately the product of the surface area of the compressed spore core *S’_core_* and the thickness *d’* of the compressed phospholipid bilayer, calculated using Eq. 6.

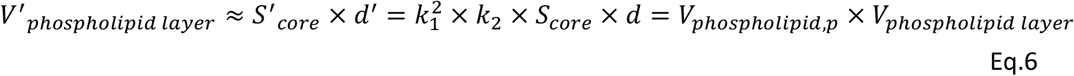

Therefore, at 30°C under HP, the relative volume of the phospholipid bilayer as a function of pressure (*V_phospholipid layer,p_*) could be calculated using Eq. 7.

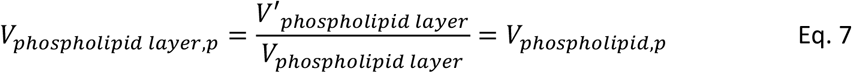

Since *V_phospholipid,p_* values from 0.1 to 100 MPa were previously calculated by Böttner et al. (69), we calculated *V_phospholipid layer,p_* values from 0.1 to 100 MPa according to their data, and presented them as the solid green line in Figure S10B. Moreover, we further calculated *V*_phospholipid layer,p_ values from 100 to 600 MPa by using predicted *V_phospholipid,p_* values according to the previous model (69), and presented them as the green dashed line in Figure S10B.

### Data processing

Unless otherwise specified, all experiments were conducted in triplicate, and the values presented are means ± standard deviations (SD) from three replicates. Statistical analyses, data processing, and graph generation were carried out using GraphPad Prism 9 software. Differences between groups were evaluated using one-way ANOVA in IBM SPSS Statistics 27, with significance levels set at p < 0.05.

## Acknowledgements

We thank Dr. Tuo Zhang (China Agricultural University, China) for valuable suggestions and discussions. We thank Prof. David Rudner (Harvard Medical School, USA) for generously gifting *B. subtilis* Δ*5* and derivative strains, which were crucial to this study. We thank Prof. Sigal Ben-Yehuda (HUJI, Israel) for gifting the germinant receptor knockout strains. This work was supported by National Natural Science Foundation of China (NSFC) (grant nos. 32522084 and 32372470), Agricultural Research Outstanding Talents of China (grant No. 13210317) awarded to Lei Rao, and 2115 Talent Development Program of China Agricultural University awarded to Lei Rao and Xiaojun Liao.

## Data availability

This paper does not report original code. The data reported in this paper will be shared by the corresponding author upon request.

